# Intra-manchette transport employs both microtubule and actin tracks

**DOI:** 10.1101/2024.10.16.618660

**Authors:** Jo H. Judernatz, Laura Pérez Pañeda, Tereza Kadavá, Albert J. R. Heck, Tzviya Zeev-Ben-Mordehai

## Abstract

The manchette is a transient microtubule based structure that plays a vital role in nuclear shaping during spermiogenesis. It comprises thousands of microtubules (MTs) that build a scaffold around the distal half of the nucleus. The manchette distributes proteins and vesicles during spermiogenesis in a process called intra-manchette transport (IMT). The current hypothesis is that IMT shares many similarities with intra-flagellar transport (IFT) and utilizes both MTs and filamentous actin (F-actin). However, IMT is still poorly understood as direct visualization of IMT complexes is missing, and the presence of F-actin has not been experimentally shown. Here, we use proteomics and cryogenic-electron tomography (cryo-ET) to identify and visualize IMT components. We find that F-actin is an integral part of the manchette with two different spatial organizations, namely bundles and single filaments, providing tracks for transport as well as having structural and mechanical roles. We further uncover that IMT on MTs is mediated by two distinct transport machineries: a dynein-mediated transport of soluble cargo and a dynein-independent transport for vesicles. Our results provide new insights into the manchette’s function as a transport scaffold, highlighting its significance for the polarization of spermatids during spermiogenesis.

## Introduction

Spermiogenesis is the post-meiotic process in which round spermatids are morphologically reorganized into motile sperm cells^1^. During spermiogenesis, the manchette assembles as a temporary microtubular scaffold around the distal half of the nucleus to aid in sperm head shaping^2–4^. Aside from the manchette function in nuclear reshaping, it provides a large cytoskeletal platform for bi-directional transport^5–7^. As such, the manchette plays a crucial role in providing a framework for the major reorganization of cellular material required to form the highly polarized sperm cell. Transport along the manchette has been termed intra-manchette transport (IMT)^7–11^. IMT has been proposed to ensure the timely delivery of macromolecules, vesicles, and mitochondria to the developing axoneme of the sperm tail involving multiprotein complexes^2,4,7,9,12^ and to redistribute nuclear material like protamines^13^.

IMT depends on the cytoplasmic microtubule (MT) motor proteins dynein and kinesin^9,10,13,14^. Cytoplasmic dynein requires the binding of dynactin to activate and tether cargo^15,16^. Once bound to dynactin, the motor domain of dyneins can undergo a mechanochemical cycle to move cargo along a microtubule^17,18^. Dyneins move along microtubules in a plus-end to minus-end direction^17^. Most kinesins, on the other hand, move along microtubules in a minus-end to plus-end direction with few exceptions^19,20^. Kinesins have four components: two identical heavy chains and two light chains. Two light chains attach to the end of the stalk, enabling cargo to bind to kinesin^17,21^. While kinesins and dynein were shown to localize to the manchette via immunofluorescence microscopy and immunogold labeling, direct observation of motor protein-mediated IMT is still lacking^10,22^.

Several other nonmotor proteins have been suggested as molecular components of the manchette and are believed to participate in IMT via direct interaction with tubulins or with dyneins and kinesins, respectively^23–25^. Subunits of intra-flagellar transport (IFT) complexes A and B were amongst the first proteins hypothesized to play a role in IMT, and all IFT genes are expressed during spermatogenesis^2^. IFT complexes are crucial for axoneme development and maintenance and form “trains” of repeating kinesin/IFT/dynein repeats moving along the developing axoneme^26^. The involvement of IFT complex B subunit IFT88 in IMT was suggested^9,27^. IFT88 mutant mice show defects in sperm head shaping and tail formation^27^. Additionally, IFT20^28^, IFT140^29^, and IFT27^30^ were shown to be localized to the spermatid manchette. However, whether IFT trains or similar structures assemble in the manchette and are involved in IMT remains unclear.

While dynein- and kinesin-mediated IMT along manchette MTs (mMT) was proposed for long distances, short-distance transport, and accurate targeting have been proposed to be carried out along actin filaments^7,8,31,32^. Actin exists as single subunits in a globular form (G-actin) and as polymerized filaments (F-actin). F-actin motor protein Myosin Va localizes to the manchette^7,8^, and actin was detected in western blots of isolated manchettes^33^. However, the presence of F-actin in the manchette remained elusive^7,34,35^. Consequently, the involvement of F-actin in the manchette remains part of an open discussion and needs clarification as concrete structural evidence for this hypothesis is lacking^36^.

Here, we surveyed the manchette proteome and used cryo-electron tomography (cryo-ET) to visualize IMT in isolated manchettes. We find that macromolecules are transported via dynein, and vesicles employ dynein-independent transport. We further show that F-actin are an integral part of the manchette forming bundles likely playing a structural and a mechanical role and single filaments providing tracks for transport.

## Results

### Vesicles are transported along manchette microtubules (mMT)

To study IMT with cryo-ET, manchettes were isolated in the presence of the MT-stabilizing agent Taxol based on a published protocol. Vesicles were highly abundant in isolated manchettes, demonstrating that they can be used to study IMT (Fig. 1).

**Fig. 1.**
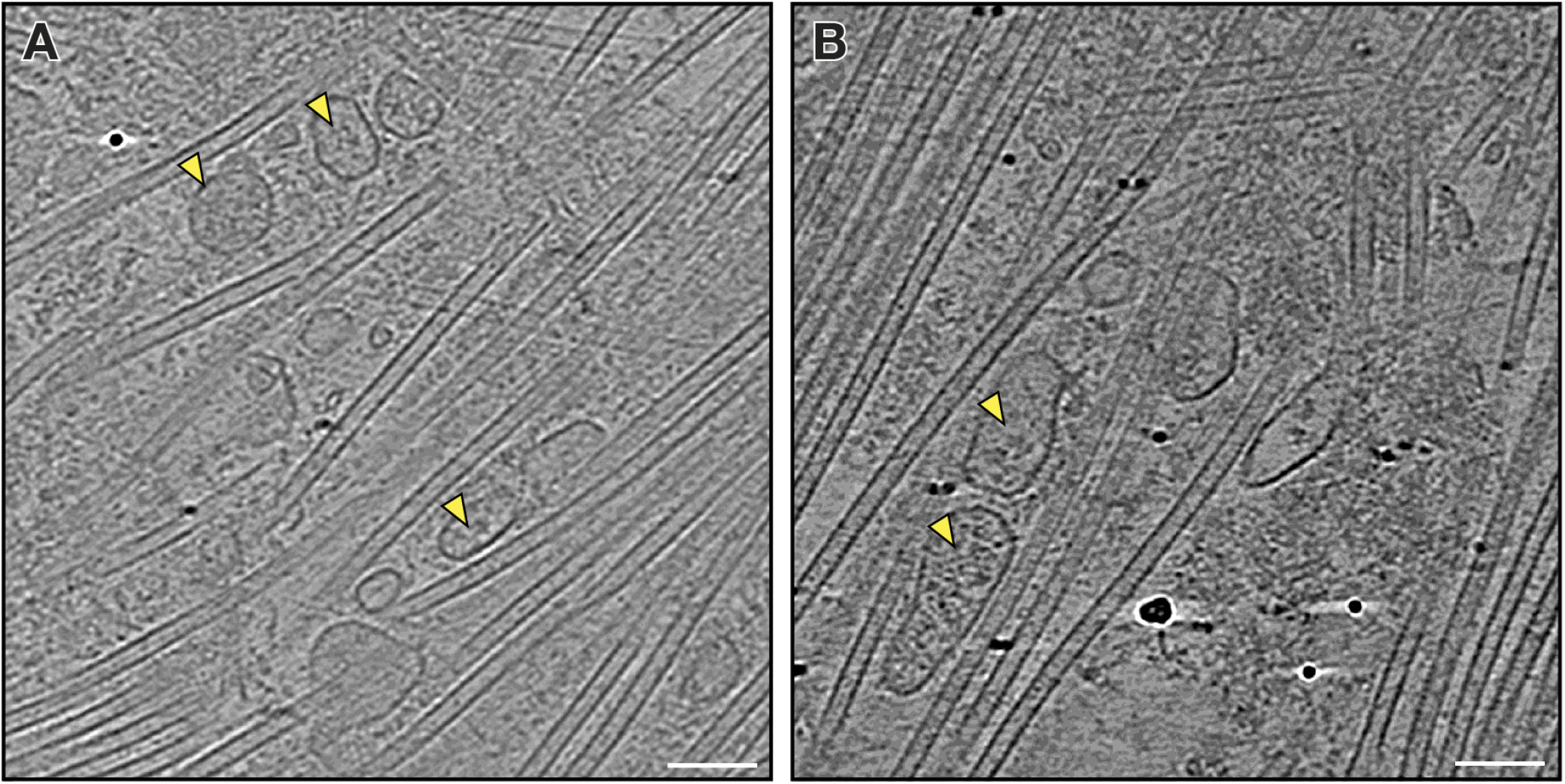
Vesicles are associated with the manchette. **(A-B)** Slices from two tomograms showing vesicles close to manchette MTs (yellow triangles). Scale bars: 100 nm.

To visualize the tethering of vesicles to mMTs, we concentrated on the manchette periphery as these regions are thinner and less crowded. We observed stick-like densities bridging some vesicles to mMT (Fig. 2A). Analysis of mMTs directionality determined by subtomogram averaging (STA) (Fig. S1) showed that the densities face towards the MT plus-end. Other vesicles were tethered to mMT with irregular densities (Fig. 2B) facing the MT minus-end. Interestingly, some vesicles were multilamellar with mushroom-shaped densities on their membranes (Fig. 2B). 3D reconstruction of these densities with STA indicated that they are ATP-synthase (Fig. 2C and Fig. S2). In agreement, we find 13 subunits of ATP-synthase in our proteomics data (Table S1). The vesicles containing ATP-synthase, however, had a different morphology than mitochondria. Yet, it is possible that the presence of detergent in the isolation buffer affected the mitochondrial integrity, and the observed vesicles with ATP-synthase are deformed mitochondria.

**Fig. 2.**
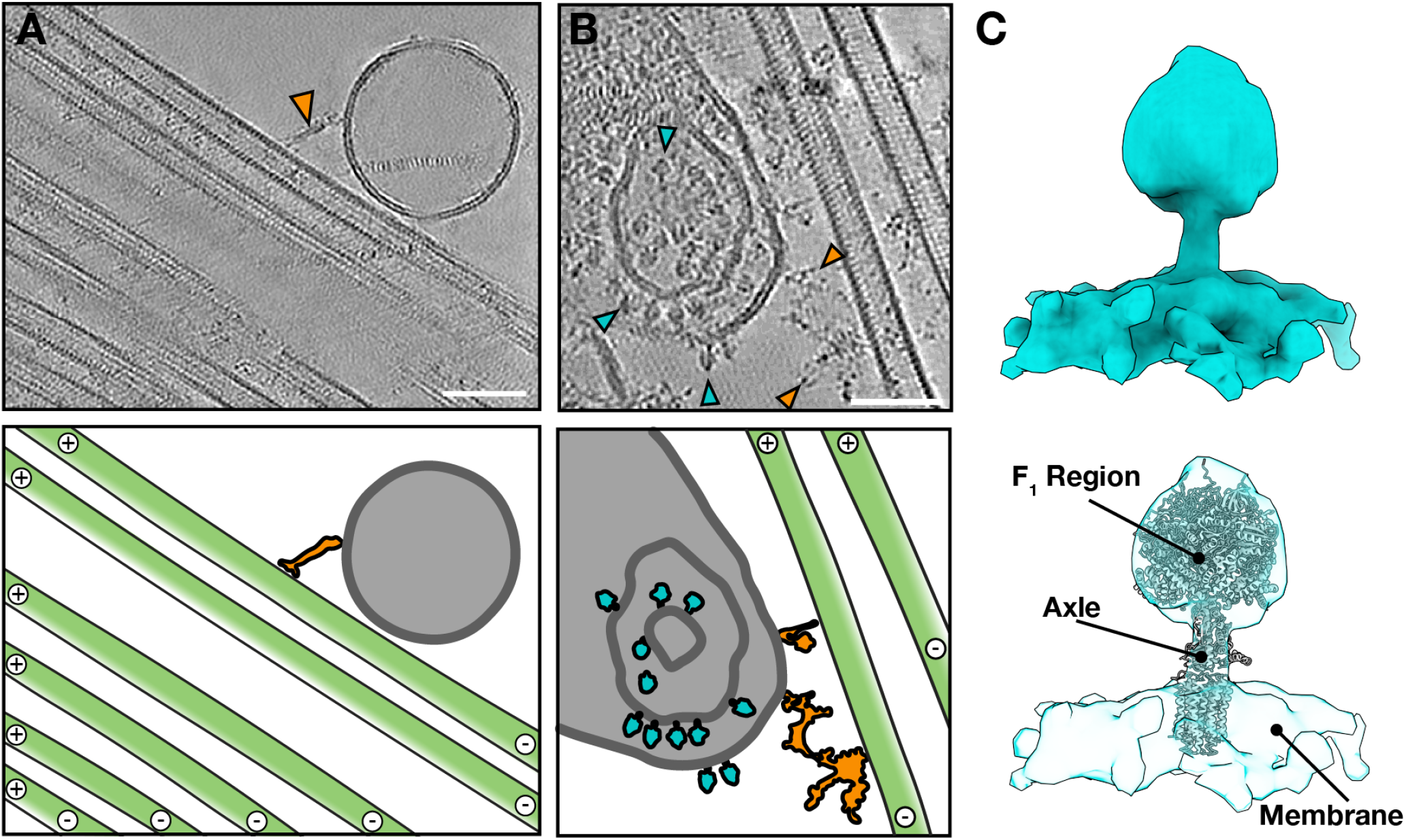
Irregular protein densities tether vesicles to manchette microtubules. **(A)** A slice through a tomogram and annotation (below) showing a vesicle (grey) connected to a microtubule (green). Scale bar: 50 nm. **(B)** A slice through a tomogram and annotation (below) of a multilaminar vesicle (light grey) with mushroom-shaped densities (turquoise) anchored to the vesicle membranes. The vesicle is tethered to a microtubule (green) via a large linker complex (orange). Scale bar: 50 nm. **(C)** A 3D reconstruction of the mushroom-shaped densities indicates that they are ATP-synthase. The F1 region and the axle of ATP synthase (PDB 6J5I) fitted well in the density (below).

### Components of the dynein transport machinery are the most abundant in the manchette

To survey the transport machinery components associated with mMTs, isolated manchettes were analyzed using bottom-up mass spectrometry (MS)-based proteomics. We classified and annotated the relative abundance of tubulins, actin, and their respective motor proteins (Fig. 3 and Table S2). As expected, the most abundant proteins were tubulin isoforms, followed by actin isoforms (Fig. 3). We found 21 kinesin superfamily members, about as abundant as subunits of cytosolic dynein complexes. Kinesins and dyneins were 10 to 100 times lower in abundance than tubulins (Fig. 3 and Table S2). The only exception is kinesin family member 27 (KIF27), which was only five times lower in abundance than tubulins (Fig. 3). KIF27 have been shown to be localized to the perinuclear ring in mouse sperm^37^ and to influence the growth dynamics of MTs^38^ but has not been implicated in transport. Members of the kinesins superfamily have diverse functions extending beyond cargo transport. Thus, comparing the relative abundance of dynein subunits with kinesins does not provide information about their potential IMT utilization. Kinesins and dynein require the formation of protein complexes to attach cargo. Kinesins depend on kinesin light chains to tether and transport cargo^39^. Likewise, dynein needs the binding of dynactins to move cargo along MTs^15^. To determine whether IMT favors the usage of either of the motor proteins, we compared the relative abundance of their respective cargo linkers. Dynactins were about 4.5 times more abundant than kinesin light chains (Fig. 3), indicating increased utilization of dyneins over kinesins in IMT.

**Fig. 3.**
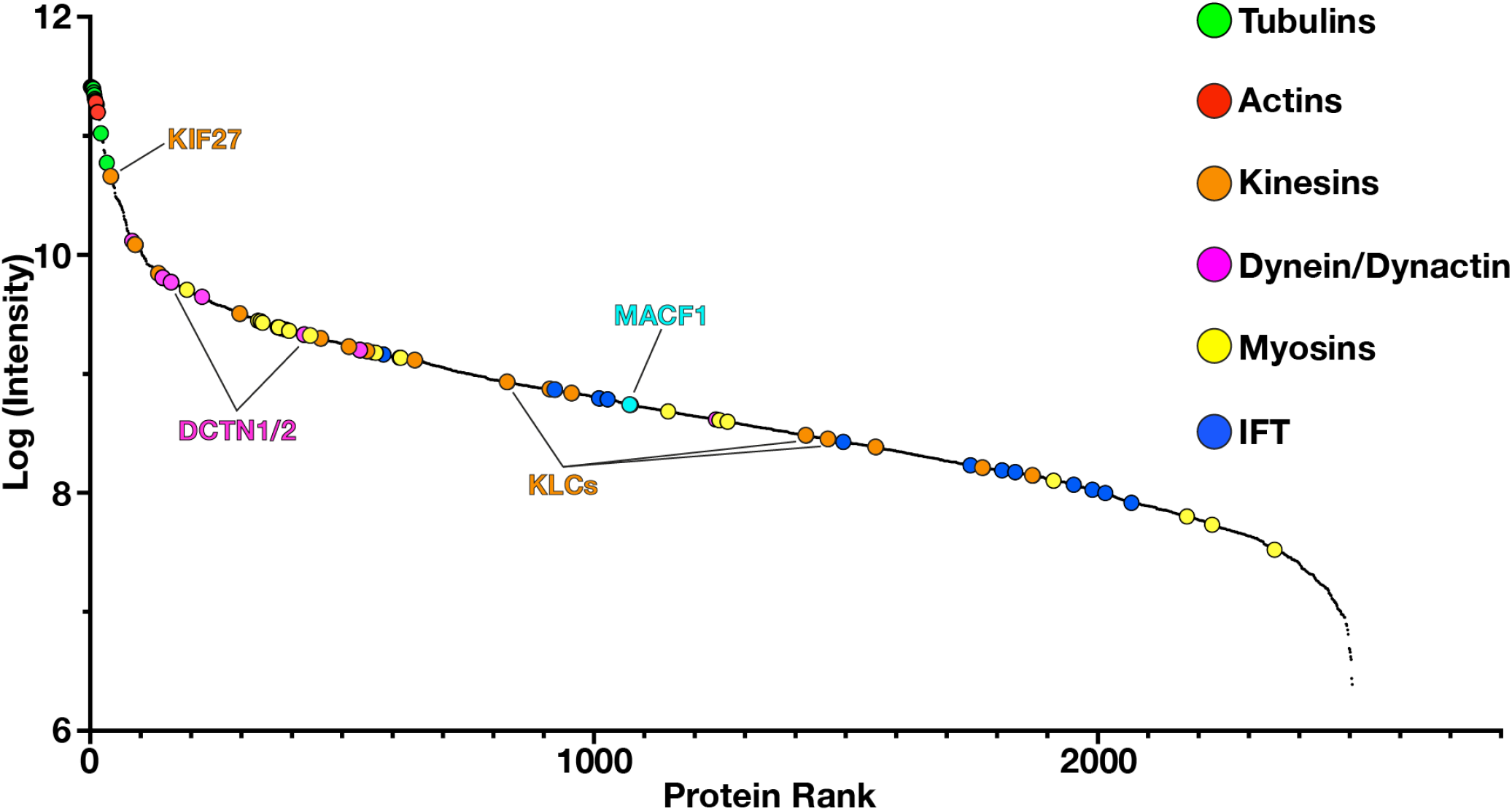
Relative abundance of transport-related components in isolated rat manchettes. Tubulins (green) and actins (red) show the highest abundance. Components of the microtubule motor proteins kinesins (orange) and dynein (pink) are similar in abundance. Components of the actin motor proteins myosins (yellow) are less abundant than dyneins and kinesins. IFT subunits (blue) show the lowest abundance.

IFT subunits are essential for proper sperm cell development^14,29,30^, and similarities between IFT and IMT have been suggested^2,4,9^. In our proteomics data, IFT sub-units were about 10 to 100 times less abundant than the microtubule motors dynein and kinesin (Fig. 3, blue circles). Indeed, we did not observe IFT trains walking along mMTs in any of our tomograms. Nonetheless, we detected the previously reported IFT subunits IFT88 and IFT20^28^ in our manchette proteomics dataset, confirming their association with the manchette. Consequently, functional roles of IFT subunits outside the IFT complex cannot be excluded from our data, and IFT proteins might be involved in binding protein cargo via different mechanisms.

### Dynein-mediated cargo along the manchette

In addition to vesicle transport along mMT, we regularly observed other cargo transport. This cargo transport is exclusively coupled to mMTs with densities showing hollow rings and slender stalks (Fig. 4A-C). 3D reconstruction of these densities had the dimensions and shape of a dynein motor domain (Fig. 4D). In each of the cases observed (48), only a single dynein motor domain was resolved (Fig. 4A-C). We found the cargo always on the side facing the plus-end mMT while the dyneins faced the minus-end, similar to other cytosolic dynein^40^. Interestingly, not all dyneins are engaged with a cargo (Fig. 4A). The cargo binding to dyneins has different sizes and shapes (Fig. 4A-C), supporting highly variable cargo in IMT needed at the caudal part of the developing spermatids, e.g., axoneme components.

**Fig. 4.**
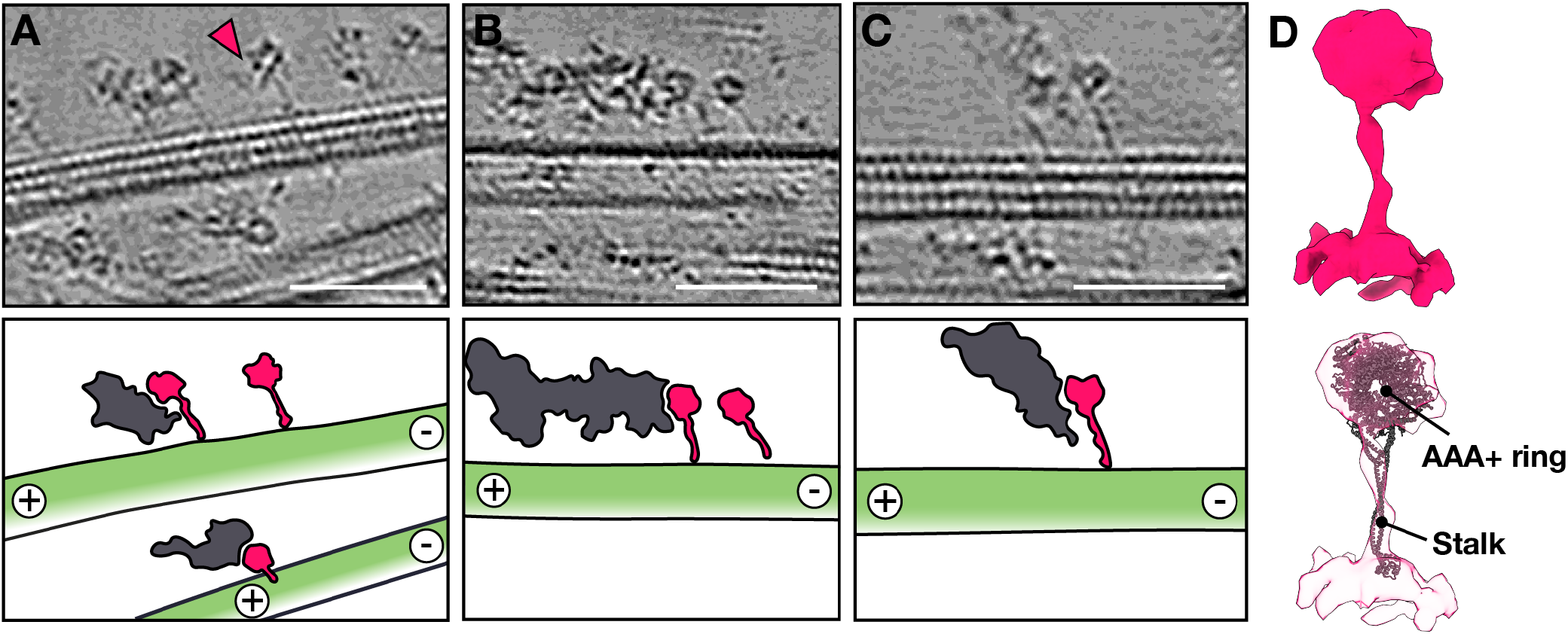
Dynein mediated transport in intra-manchette transport. **(A-C)** Slices through tomograms and annotations showing transport proteins (pink) carrying cargo (grey) towards the minus-end of mMTs (green). Pink triangles indicate dynein motor domain without attached cargo. Scale bars: 50 nm. **(D)** 3D reconstruction of the transport proteins with a crystal structure of dynein II motor domain (PDB: 4RH7) fitted into the map (below).

### Filamentous-actin are an integral part of the manchette

In addition to MTs, we consistently observed non-MT filaments as part of the manchette (Fig. 5 and 6). 3D reconstruction of these filaments revealed dimensions and a helical twist of F-actin (Fig. 5B). In our proteomics data, actin isoforms alpha-actin-2 and beta- and gamma-actin were highly abundant, supporting the assignment of F-actin (Fig. 3 and Table S2).

**Fig. 5.**
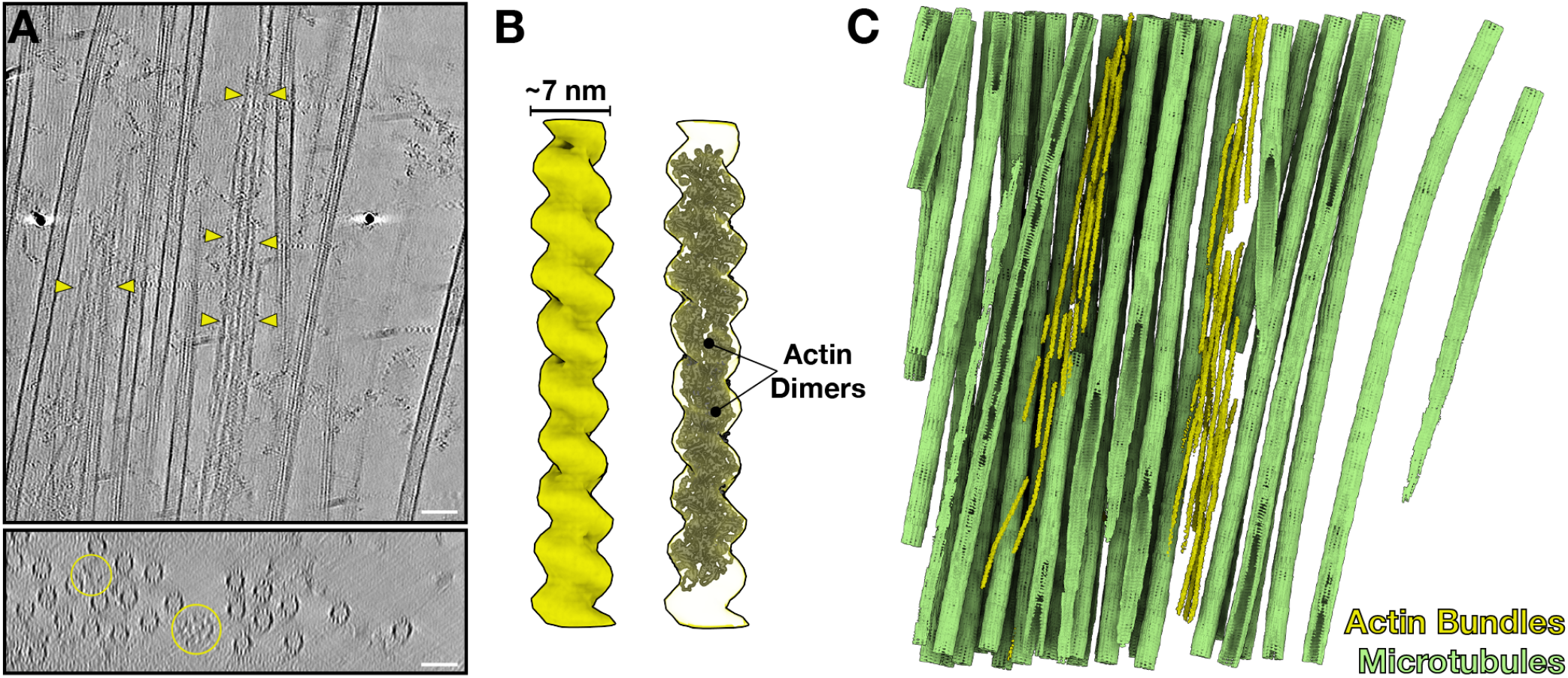
F-actin forms bundles in the manchette. **(A)** Orthogonal slices from a tomogram showing non-MT filaments (yellow arrowheads and yellow circles) forming bundles. Scale bar: 50 nm. **(B)** 3D reconstruction and fitting of an F-actin structure (PDB:7BT7). **(C)** Segmentation of the tomogram in A with F-actin (yellow) bundles running parallel in between mMTs (green).

**Fig. 6.**
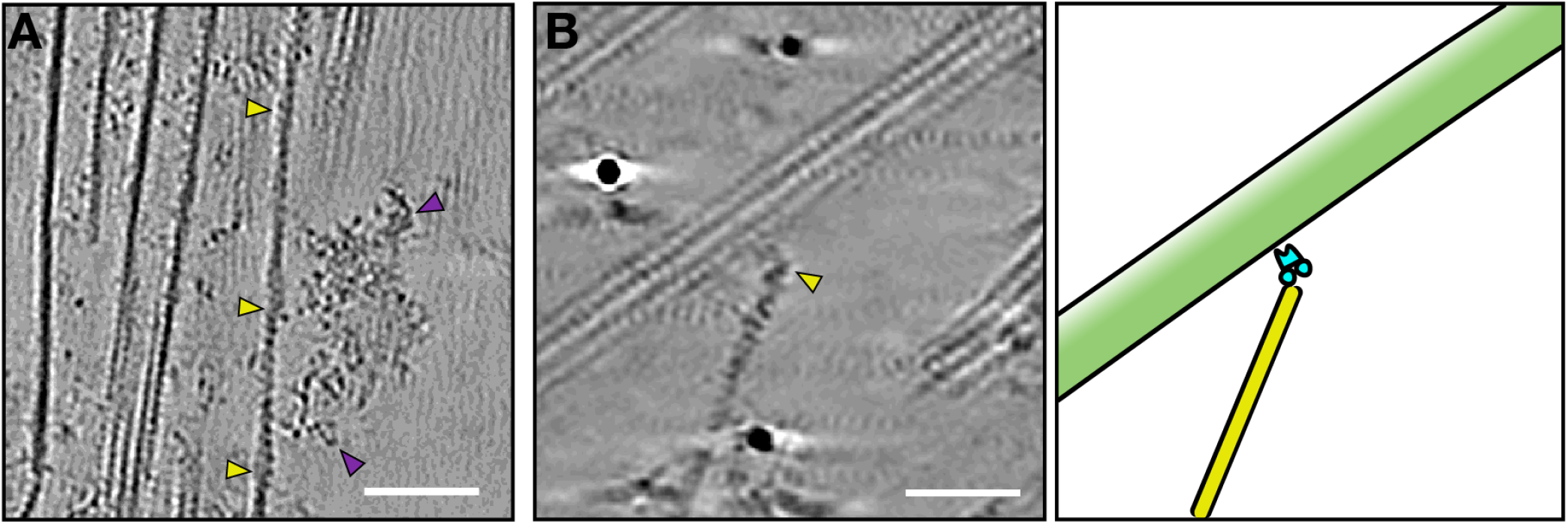
F-actin singlets provide tracks for transport. **(A)** A slice from a tomogram with a single actin filament (yellow arrowheads) bound by a large cargo density (violet arrowheads). Scale bar: 50 nm. **(B)** A slice from a tomogram with an actin singlet interacting with mMT through a linker protein. Annotation on the right. MT, green; Actin, yellow; Linker protein, teal. Scale bar: 50 nm.

F-actin formed organized bundles running parallel to mMTs (Fig. 5). Bundles comprised two to ten single actin filaments and were arranged tightly between mMTs (Fig. 5). The actin filaments were shorter than mMTs and often seemed too short for meaningful use in cargo transport. It is, however, possible that F-actin were shorter in our isolated manchette due to sample preparation, as no F-actin stabilizer was added. F-actin bundles formation requires crosslinking proteins^41^ and in our proteomics we detected the actin cross-linking proteins espin, a-actinin, and fascin (Table S2). While α-actinin and fascin showed a relatively low abundance of 100 to 1000 times less than actin, espin was only four times lower in abundance compared to actin. Nevertheless, we have not observed densities between F-actin. In addition to the F-actin bundles, we found single F-actin filaments (or singlets) close to mMTs (Fig. 6A,B). Unlike F-actin bundles that were naked, F-actin singlets occasionally had large protein complexes attached (Fig. 6), potentially motor proteins with cargo. In agreement with a potential role for actin in transport, myosins were only slightly lower in abundance than kinesins and dynein subunits in the proteomics data (Table S2). In one instance, we observed a link between an actin singlet and a mMT (Fig. 6B). In the proteomics data we find MT actin-crosslinking factor 1 (MACF1) (Fig. 3) as a possible candidate for this linker. The two organizations of F-actin (bundles and singlets) in the manchette hint towards a dual role for actin. Singlets could function in cargo transport, while bundles might take a structural or mechanical role in the manchette.

## Discussion

The shaping of sperm during spermiogenesis involves extensive morphological remodeling that requires the relocation of molecular building blocks within the spermatid. The manchette has been proposed to play a crucial role as a platform for bi-directional cargo transport, ensuring the timely delivery of macromolecules in the spermatid to form the highly polarized sperm cells^9^. Here, we have shown that dyneins play an active role in IMT (Fig. 4). Our proteomics data further suggests that IMT via dyneins is predominant. Dynein motors walk along MTs only in a plus-to minus end direction^42^. In the manchette, the MT plus-ends are organized unidirectionally, with the plus-ends facing the perinuclear ring and minus-ends leading toward the developing axoneme^2,43^. This unidirectionality of dynein-mediated IMT suggests that the cargo we observe could be components of the axoneme and neck region of the spermatid.

Our data further indicates that the transport of vesicles along mMTs does not depend on dyneins but rather a different transport machinery^3^. Vesicles are associated with mMTs through stick-like densities (Fig. 2), as observed with classical electron microscopy^3^. Aside from dyneins, our proteomics showed the presence of multiple kinesin-like family members, some of which have previously been detected, in the manchette (KIF2A, KIF3B, KIF27, and KIF5C)^37,44,45,46^. Kinesins are mostly plus-end directed motors; however, some kinesins with MT minus-end directionality have been found^19^. Kinesins are likely involved in the IMT of vesicles to convey polarization of the spermatid. However, kinesins show diverse functionalities and further studies are required to understand the exact nature of kinesin-mediated IMT. We did not observe densities resembling assembled IFT trains in our tomograms and did not find all subunits of the IFT complexes in the proteomics of the purified manchettes. Hence, different functionalities of particular IFT complex subunits outside of known IFT should be considered during IMT and require further investigation.

We find that F-actin are an integral part of the manchette and are present as parallel bundles as well as singlets between mMTs (Fig. 5, Fig. 6). These two forms of F-actin indicate a dual role for F-actin in the manchette: a role in transport and a mechanical role. F-actin bundles in other cells are often associated with mechanical functions, e.g., generating forces relevant to cellular processes like cell migration and cell division. In the manchette, the F-actin bundles might apply contractile forces needed for nuclear shaping. F-actin bundles are formed by crosslinking proteins^41,47^, and our proteomics data show the crosslinking protein espin in high abundance. F-actin singlets seem to function in cargo transport (Fig. 6). F-actin have previously been proposed to facilitate accurate short-distance IMT. We observed interactions between F-actin singlets and mMTs (Fig. 6) that could potentially facilitate ‘ track’ changes between MTs and F-actin. A good candidate for the protein linking mMT and F-actin is MACF1, which we find in our proteomics dataset (Fig. 3). The rare observation of linkages between F-actin and manchette MTs made structural analysis impossible, but immunofluorescence or mouse KO experiments could unveil the function of MACF1 in the manchette.

In conclusion, our data show that MT and F-actin bundles are both constituents of the manchette. They further reveal that IMT employs different transport machineries on mMT, with macromolecules using a dynein mediated transport and potentially kinesin for vesicle transport. Transport on MT tracks is augmented by transport on F-actin singlets.

## Acknowledgements

We thank the Instantie voor Dierenwelzijn (IvD) Utrecht and Ate Bijlsma for providing and sacrificing the rats. Cryo-ET data were collected at the Utrecht University Electron Microscopy Centre and at the Netherlands Centre for Electron Nanoscopy (NeCEN). We thank Dr. M. Vanevic for computational support at Utrecht University, and acknowledge Dr. S.C. Howes, M. Bergmeijer, Ingr. C. Schneijdenberg, and J. Meeldijk for management and maintenance of the Utrecht University EM Centre. Willem Noteborn for help with data collection, Miguel Leung for help with cryo-ET data processing. This work benefited from the Netherlands Electron Microscopy Infrastructure (NEMI), project number 184.034.014 of the National Roadmap for Large-Scale Research Infrastructure of the Dutch Research Council (NWO). The proteomics analysis was supported by the NWO through funding for the Netherlands Proteomics Centre through the X-omics Road Map program (project 184.034.019). T.Z. was funded by the European Research Council (ERC-2022-COG project 101088673).

## Author Contributions

J.H.J. and T.Z.B.M. conceived the project, performed experiments, collected and analyzed the data, and wrote the paper. L.P.P., T.K., and A.J.R.H. collected and analyzed mass spectrometry data. T.Z.B.M. acquired funding and coordinated the project. All authors reviewed and edited the paper.

## Data Availability

All LC-MS/MS data have been deposited to the ProteomeXchange Consortium via the PRIDE^58^ partner repository with the dataset identifier: PXD055902

## Materials and Methods

### Manchette isolation

Isolation of manchettes from rat testes was based on a previously published protocol^33^. Briefly, testes were prepared from the abdomen of freshly asphyxiated rats (Wistar, Crl:CD(SD), RJHan:WI, Lister Hooded). Fat tissue and epididymis were carefully removed using sharp scissors. The testes were then transferred to a culture dish containing PBS on ice, and the tunica albuginea was removed using tweezers.The seminiferous tubules were briefly washed and then transferred into 10 ml of microtubule-stabilizing buffer (MSB: 25 mM HEPES, 2.5 mM MgSO4, 2.5 mM EGTA, 15 mM KCl, 5 mM DTT, 0.1 mM GTP, 20 µM Taxol, 1 % Triton X-100, 1 Tablet of cOmplete mini-Protease Inhibitor). Next, the seminiferous tubules were chopped using a razorblade and then repeatedly pipetted with a 1 ml pipet to break up the tissue and separate the manchettes. The suspension was filtered through 100 µm, 30 µm, and 10 µm mesh-sized cell strainers and collected in a 50 ml Falcon tube. The cell suspension was further filtered through 7.5 g of 212-300 µm glass beads packed on a 20 mL syringe and prewashed with ice-cold PBS. After filtration, 6 mL of the suspension was added to 44 mL cold 2.5 M sucrose. The resulting 50 mL suspension was split equally into two open-top thin wall ultra-centrifugation tubes (Beckmann Coulter 344058). The mixture was layered with 6 mL of cold 2.05 M sucrose followed by 6 mL of cold 1 M sucrose and centrifuged at 85000 *x g* for 110 min at 4°C using a SW32 rotor in a Beckman Coulter Optima™ XPN-80 ultracentrifuge. Approximately 3 mL of intact manchettes were harvested from each tube at the interface between the 1 M and 2.05 M sucrose and transferred into 12 mL cold PBS. The mixture was then centrifuged at 1000 *x g* for 30 min at 4°C, the supernatant was discarded, and the pellet was resuspended in 500 µL PBS. The manchette-containing pellets were washed three times by resuspension in cold PBS and finally used for further analysis.

### Cryo-ET sample preparation and data collection

Isolated manchettes were mixed with BSA-conjugated gold beads (Aurion) in a 4:1 ratio, and 3.5-4 µL of the mixture were applied to glow-discharged Quantifoil R 2/1 200-mesh holey carbon grids. The grids were blotted from the back for 3-4 s with Whatman 1 filter paper on a manual plunger (MPI Martinsried, Germany). Grids were plunged into liquid ethane cooled to liquid nitrogen temperatures. Cryo-ET data of manchettes was collected on a 200 kV Talos Arctica (Thermo Fisher Scientific) with a pixel size of 2.17 Å or on a 300 kV Titan Krios with a pixel size of 6.32 Å. A total of 2 datasets were collected from a total of 4 grids from 2 separate manchette preparations. Tilt series were collected using SerialEM^48^ at a target defocus of between -2 and -4 µm. Tilt series were typically recorded using strict dose-symmetric schemes, spanning ± 51° in 3° increments, with the total dose limited to around 100 e-/Å^2^.

### Tomogram reconstruction

Motion between individual frames was corrected using MotionCor2 1.2.1^49^. Tomogram reconstruction was performed in either IMOD or AreTomo^50,51^. In IMOD, four-times binned tomograms were reconstructed using weighted back-projection, with a SIRT-like filter applied for visualization/segmentation and CTF-corrected using IMODs ctfphaseflip function for subtomogram averaging in PEET 1.13.0^52,53^. In AreTomo, four-time binned tomograms were reconstructed using weighted-back projection. AreTomo reconstructed tomograms were denoised with CryoCare^54^ for visualization purposes.

### Determination of MT directionality

Individual MTs were manually traced in IMOD, and model points were imported to the software package cylindra^55^. In cylindra, the protofilament skew and numbers were determined for each MT using local averages. A clockwise protofilament skew is aminus-to plus-end direction. A counterclockwise skew is a plus-to minus-end direction.

### Subtomogram averaging of dynein motor domains

Individual particles were manually picked in IMOD using low-pass filtered sub-tilt tomograms four times and binned to a final pixel size of 8.68 Å. Two points indicating the top and bottom of the dynein motor domain relative to the microtubule were picked per particle. A total of 49 particles were picked from four tomograms and used for STA in PEET 1.13.0^52,53^. Initial particle orientation and rotation axes of particles were generated relative to the MT via the SpikeInit function of PEET and used as initial information for the alignment. Resolution was estimated using the Fourier shell correlation (FSC) at a cut-off of 0.5 (Fig. S3).

### Subtomogram averaging of ATP-synthase

Individual particles were manually picked in IMOD using low pass filtered sub-tilt tomograms four times binned to a final pixel size of 8.68 Å. Two points indicating the top and bottom relative to the membrane were picked per particle. A total of 69 particles were picked from two tomograms and used for STA in PEET 1.13.0. Initial particle orientation and rotation axes of particles were generated via the SpikeInit function of PEET and used as initial information for the alignment. Resolution was estimated using the Fourier shell correlation (FSC) at a cut-off of 0.5 (Fig. S3).

### Subtomogram averaging of actin filaments

Individual actin filaments were manually traced in IMOD, and model points were added every 3.6 nm using addModPts. Subtomogram averaging with missing wedge compensation was performed using PEET 1.13.0. Alignments were generally performed first on four-times binned data. Subtomograms of approximately 16 nm × 16 nm × 16 nm were computationally aligned and averaged in steps of tighter angular and translational search ranges. A tight cylindrical mask was used during alignment to exclude signal from neighboring actin filaments. A total of 928 particles from two tomograms contributed to the final average with a resolution of 30 Å. Resolution was estimated using the Fourier shell correlation (FSC) at a cut-off of 0.5 (Fig. S3).

### Proteomics sample preparation

Single-pot solid-phase-enhanced sample preparation (SP3)^56^ was used for sample processing for proteomic analysis. Samples were lysed with 20 µL of lysis buffer (1 % (w/v) sodium dodecyl sulfate, 100 mM HEPES, pH 8, 1 % (v/v) EDTA-free complete protease inhibitor (Roche)) and sonicated in the Bioruptor (Diagenode) for 15 cycles of 15s on/off intervals. Benzonase (Purity >90 %, Millipore) was added to 1 % (v/v) to remove RNA or DNA contaminants, and the sample was incubated at 37°C for 15 min. The lysates were centrifuged for 10 min at 18,000 *x g* at 4°C, and the supernatant was collected. A bicinchoninic acid assay was performed to estimate protein concentration. Each sample was split into three equal volumes and diluted with lysis buffer.

Samples were reduced and alkylated for 5 min at 95 °C by adding tris(2-carboxyethyl)phosphine and chloroacetamide to a final concentration of 40 and 10 mM, respectively. Pre-washed SeraMag paramagnetic beads (Thermo Fisher Scientific) were added at a 1:10 bead-to-protein ratio, and protein binding was induced by adding ethanol to 75 % (v/v). Samples were incubated for 20 min at 1000 rpm at room temperature. After incubation, the beads were rinsed in the magnetic rack twice with 200 µL of 80 % (v/v) ethanol and once with 200 µL of 100 % (v/v) acetonitrile. The proteins were resuspended in 100mM ammonium bicarbonate, and samples were sonicated in a water bath for 2 min. Protein digestion was achieved by adding trypsin and LysC at 1:25 and 1:75 enzyme-to-protein ratios, respectively. Samples were incubated at 37 °C overnight (ca. 16h) at 1000 rpm in a thermo shaker.

Samples were centrifuged for 2 min at 18,000 *x g* and acidified with 5 % (v/v) trifluoroacetic acid. The beads were immobilized in the magnetic rack, and the peptide solution was recovered. Finally, sample clean-up was performed using an Oasis HLB 96-well µElution Plate (Waters). Peptides were dried in a SpeedVac concentrator and re-suspended in 0.1 % formic acid before liquid chromatography with tandem mass spectrometry ((LC)-MS/MS) analysis.

### LC-MS/MS analysis

Peptide samples were analyzed using a nanoUltimate 3000 UHPLC (Thermo Fisher Scientific) coupled to an Orbitrap Exploris 480 mass spectrometer (Thermo Fischer Scientific). The sample (1 µg) was injected at 3 µL/min for 1 min into a trap column (Acclaim Pepmap 100 C18, 5 mm x 0.3 mm, 5 µm, Thermo Fisher Scientific). The peptide separation was performed in a 50 cm long analytical column with a 75 µm inner diameter packed in house with C18 beads (Reprosil C18, 1.9 µm) at a column temperature of 32°C. Peptides were eluted at 300 nL/min with a total run time of 90 min. Gradient separation on the analytical column: 9 % B (80 % acetonitrile and 0.1 % formic acid) for 1 min, from 9 % to 13 % B in 1min, from 13 % to 44 % in 65 min, from 44 % to 55 % in 5min, from 55 % to 99 % in 3 min, 99 %B for 5 min and 9 % B for 10 min. The MS acquisition method was a data-dependent acquisition mode using the Orbitrap analyzer at 60K mass resolution in the scan range 375-1600 m/z, with automatic maximum injection time and a standard AGC target. Ions subjected to MS2 were filtered based on dynamic exclusion with a mass tolerance of 10 ppm. Peptides were fragmented with HCD of 28 % and mass resolution of 15K.

### MS Data analysis

The raw files were analyzed with FragPipe version 20.0 ^57^. Data were searched against the reviewed and unreviewed Uniprot rat proteome (Uniprot Proteome ID: UP000002494, downloaded November 2023), and known contaminants. The precursor and the fragment mass tolerance were set to 20 ppm. Full tryptic digestion with a maximum of three missed cleavages was allowed. Carbamidomethylation (Cys) was set as fixed modification, and oxidation (Met) and acetylation (protein N-terminus) were set as dynamic modification. The false discovery rate (FDR) was set to 1 %. Further data analysis was performed in the Rstudio programming language environment. Protein intensities were extracted from FragPipe output. The data was filtered, removing proteins without quantitative values, with only shared peptides, and contaminants. Only proteins quantified in 2 out of 3 replicates were further used for the analysis.

---

**Fig. S1.**
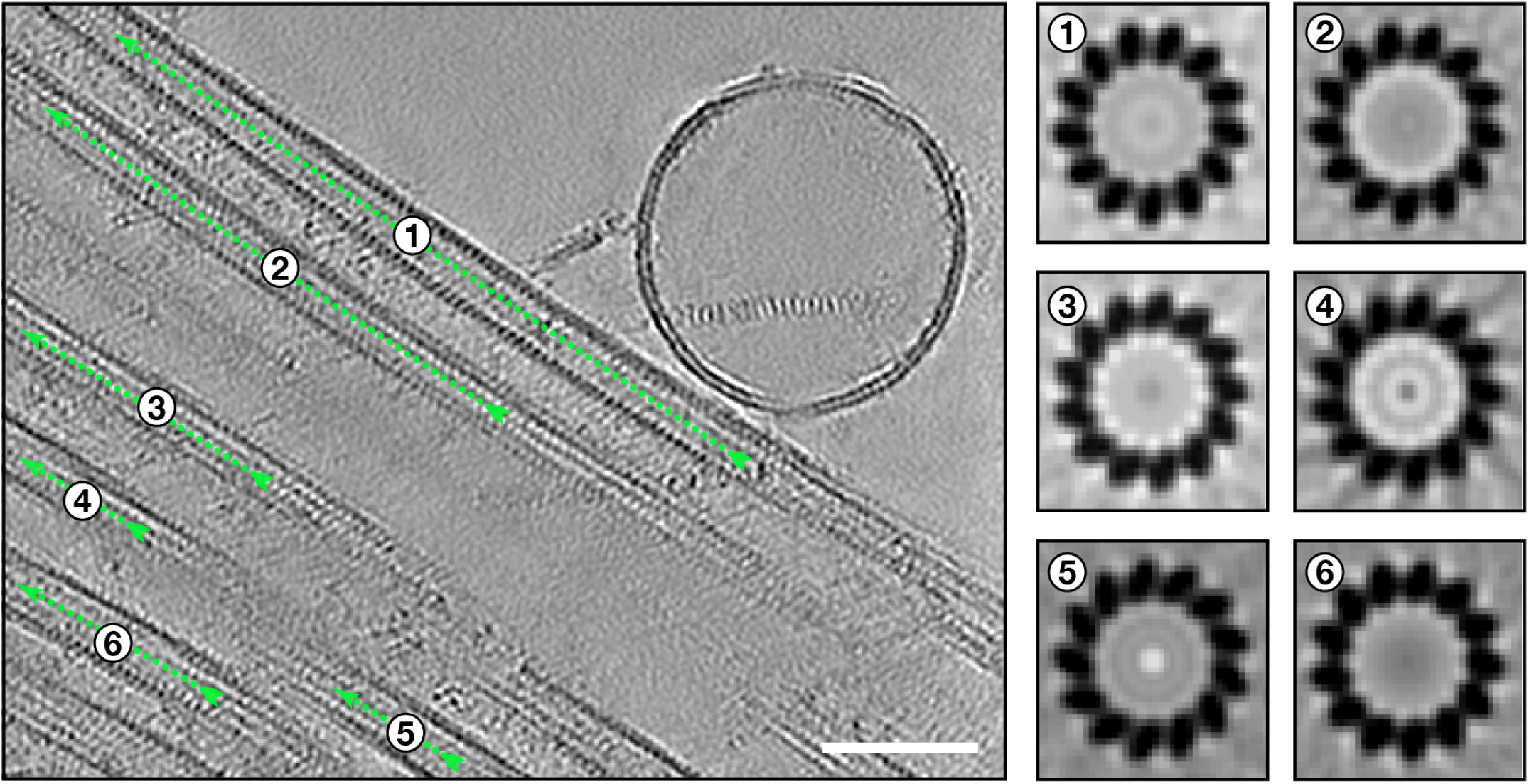
MT protofilament slew indicates its directionality. Green arrows show the direction of view. A clockwise protofilament (PF) slew is a view from the minus-to plus-end direction. Scale bar: 50 nm.

**Fig. S2.**
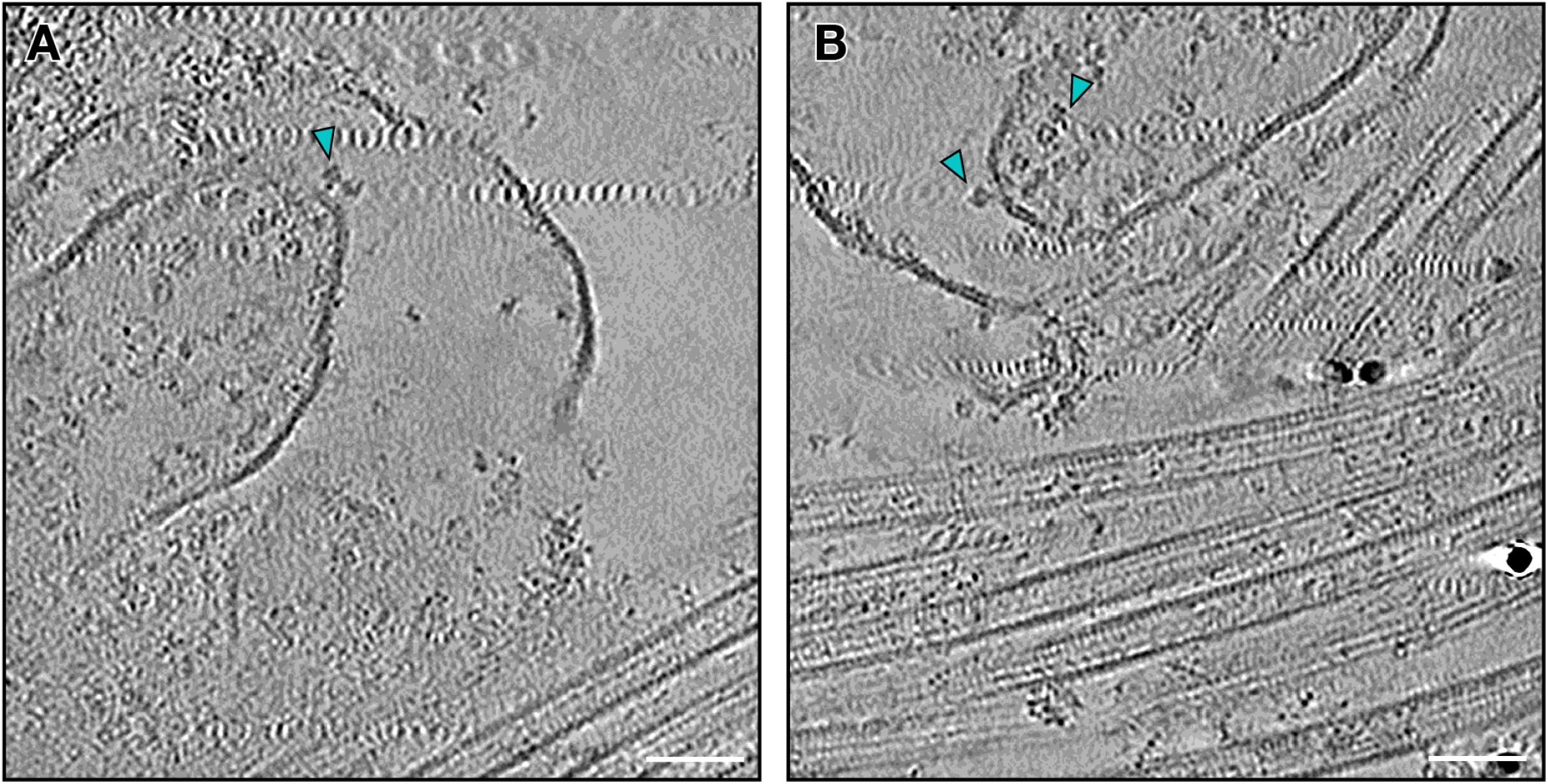
Vesicles with ATP-synthase on their membranes (cyan triangles). Scale bar: 50 nm.

**Fig. S3.**
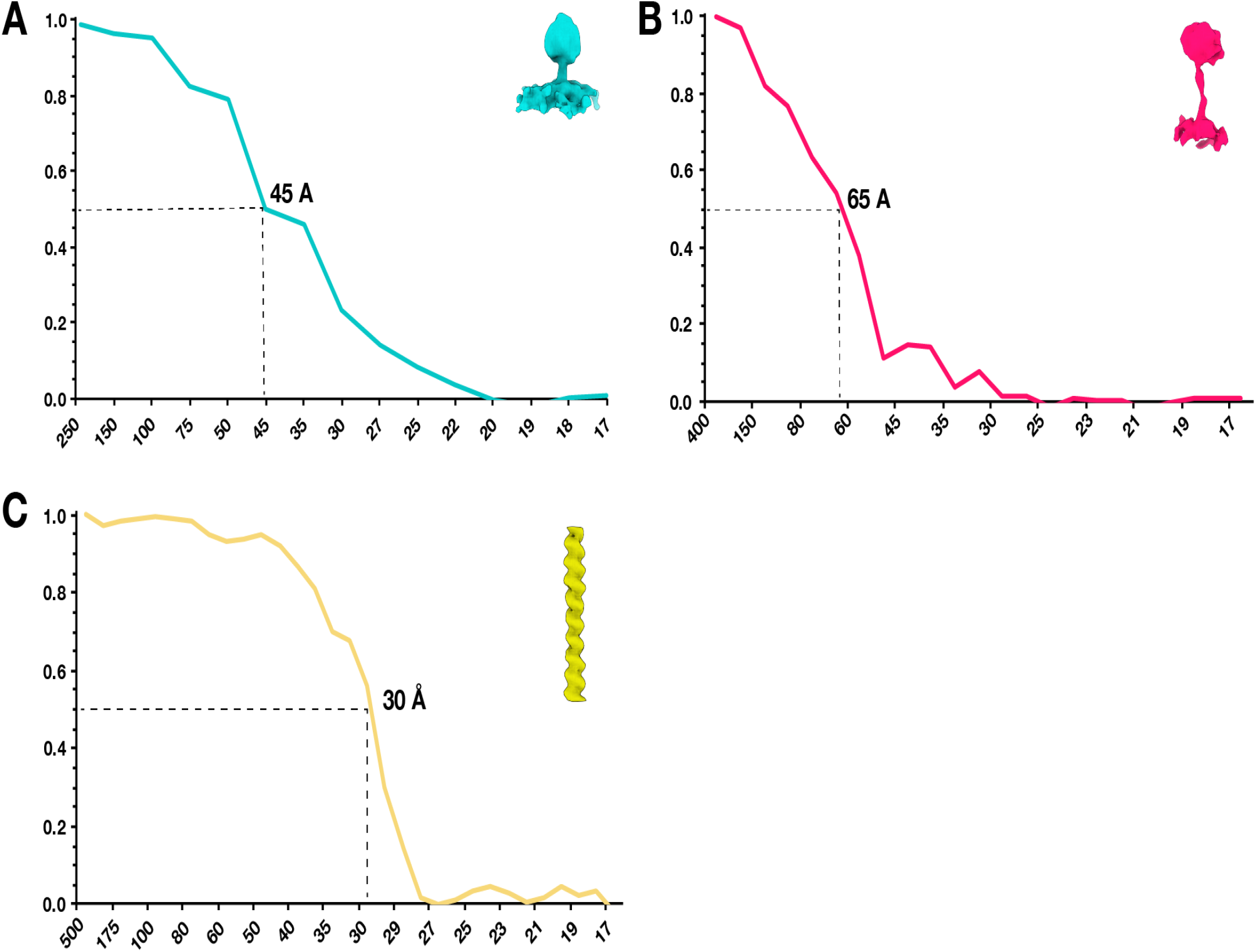
Fourier shell correlations of subtomogram averages. **(A)** FSC curve of subtomogram average of ATP-synthase. **(B)** FSC curve of subtomogram average of dynein motor domains. **(C)** FSC curve of subtomogram average of F-actin. Resolutions were estimated based on a cutoff at 0.5.

**Table S1.** Relative abundance of subunits of ATP synthase identified in proteomics data of isolated rat manchettes.

| **Protein ID** | **Gene name** | **Intensity** | **Log Intensity** | **MW(kDa)** |
| --- | --- | --- | --- | --- |
| P15999 | Atp5f1a | 3,71E+09 | 9,57 | 59,754 |
| P19511 | Atp5f | 1,03E+09 | 9,01 | 28,869 |
| P35435 | Atp5f1c | 9,40E+08 | 8,97 | 30,191 |
| P31399 | Atp5h | 8,81E+08 | 8,94 | 18,763 |
| D3Z9R8 | Atp5mpl | 8,25E+08 | 8,92 | 6,914 |
| Q06647 | Atp5po | 6,13E+08 | 8,79 | 23,398 |
| P35434 | Atp5f1d | 5,54E+08 | 8,74 | 17,595 |
| P29419 | Atp5me | 3,36E+08 | 8,53 | 8,255 |
| D3ZAF6 | Atp5mf | 3,12E+08 | 8,49 | 10,452 |
| Q9JJW3 | Atp5mk | 2,67E+08 | 8,43 | 6,408 |
| Q6PDU7 | Atp5mg | 1,86E+08 | 8,27 | 11,433 |
| P21571 | Atp5pf | 1,63E+08 | 8,21 | 12,494 |

**Table S2.**
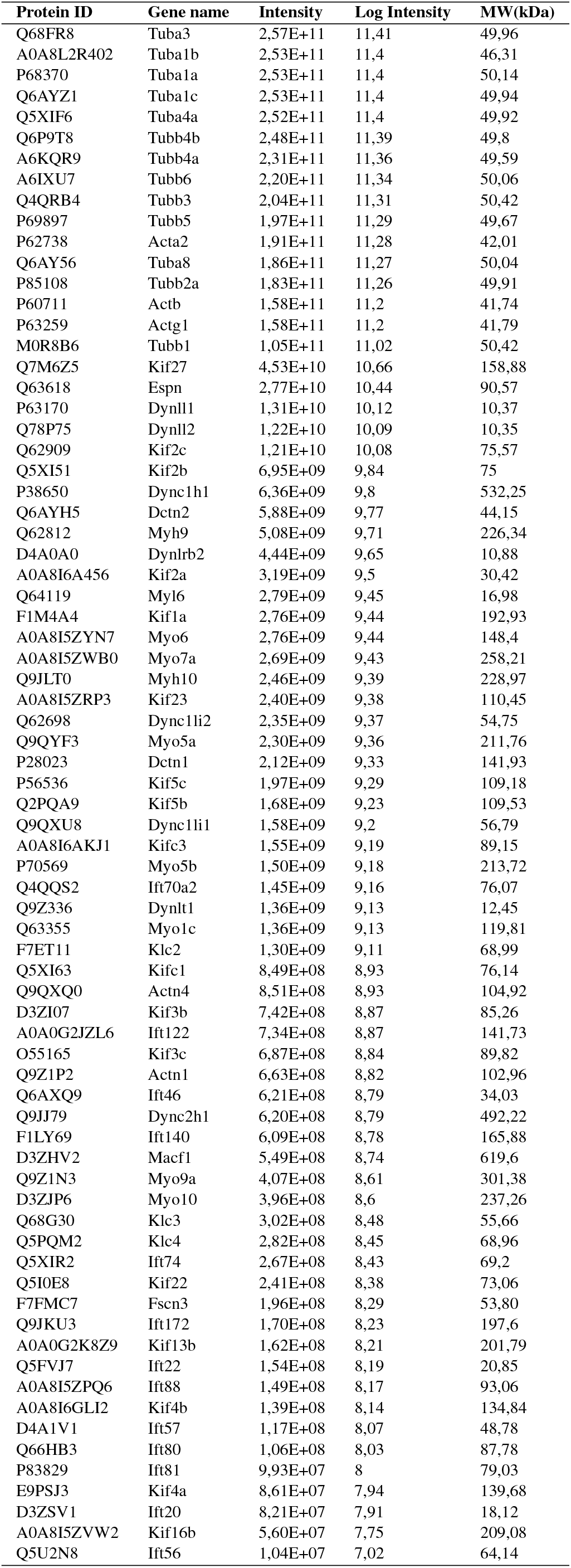
Relative abundance of tubulins, actins, and interacting proteins found in proteomics of isolated rat manchettes.

